# A multi-structural finite element model to simulate atomic force microscopy nanoindentation of single cells

**DOI:** 10.1101/2019.12.17.879114

**Authors:** Stefania Marcotti, Gwendolen C Reilly, Damien Lacroix

**Affiliations:** Insigneo Institute for *in silico* Medicine, Sheffield, UK; Dept. of Mechanical Engineering, University of Sheffield, UK; Randall Centre for Cell and Molecular Biophysics, King’s College London, UK; Dept. of Materials Science and Engineering, University of Sheffield, UK

## Abstract

Single cell mechanical properties represent an increasingly studied descriptor for health and disease. Atomic force microscopy (AFM) has been widely used to measure single cell stiffness, despite its experimental limitations. The development of a computational framework to simulate AFM nanoindentation experiments could be a valuable tool to complement experimental findings. A single cell multi-structural finite element model was designed to this aim by using confocal images of bone cells, comprised of the cell nucleus, cytoplasm and actin cytoskeleton. The computational cell stiffness values were in the range of experimental values acquired on the same cells for nanoindentation of the cell nucleus and periphery, despite showing higher stiffness for the nucleus than for the periphery, oppositely to the average experimental findings. These results suggest it would be of interest to model different single cells with known experimental effective moduli to evaluate the ability of the computational models to replicate experimental results.

## Introduction

Single cell mechanical properties are increasingly regarded as a valuable descriptor of cell physiological and pathological conditions [7]. These properties have been tested experimentally with atomic force microscopy (AFM) force spectroscopy [3], a technique allowing to perform controlled nanoindentation of biological samples [7, 12]. The determination of the effective modulus of such samples is, however, prone to various factors leading to experimental error, such as the experimental set-up, the testing conditions and the performed data analysis [3, 7–9]. The methodological variability factors can be reduced by careful optimisation of the experimental protocols. However, it is not always straightforward to perform accurate sample selection due to the time- and user-intensive nature of the AFM experiments [3]. It would be, therefore, of interest to simulate AFM force spectroscopy experiments within a computational framework, as this would allow for a controlled virtual environment exempt from experimental limitations. The use of confocal microscopy based finite element (FE) models with an accurate representation of intracellular structures can strongly support experimental work on cell mechanics [15, 17]. These models, in fact, allow for realistic representation and quantification of mechanical stimuli acting on individual cells. Moreover, numerical simulations including different cell components provide insights into the contribution of individual elements to whole-cell mechanics [1] and on the distribution of forces for structural stability [11]. An image-based single cell model was created to simulate AFM nanoindentation experiments. The model comprised the nucleus, the cytoplasm and the actin cytoskeleton. The latter was modelled as actin bundles resisting tensional forces [1,11, 15, 18] and actin cortex responding to compression [1, 18]. The use of confocal stack images of a representative cell of a bone cell line allowed for faithful representation of the cell shape and of the arrangement of the actin bundles. AFM indentation was simulated in different cell regions to investigate the effect of the compression of multiple intracellular components with different mechanical properties. AFM force spectroscopy data obtained on the same cells were used for comparison in terms of cell stiffness.

## Materials and Methods

### Cells

Pre-osteocyte MLO-A5 cells were used [6], kindly donated by Prof. Lynda Bonewald (University of Missouri). Confocal Z-stack imaging of cell samples stained for actin with Phalloidin–Tetramethyl rhodamine B isothiocyanate (TRITC) and for nuclei with 4’,6-diamidino-2-phenylindole (DAPI) was performed (Fig. 1B) at the Microscopy Core Facility (University of Sheffield).

**Figure 1.**
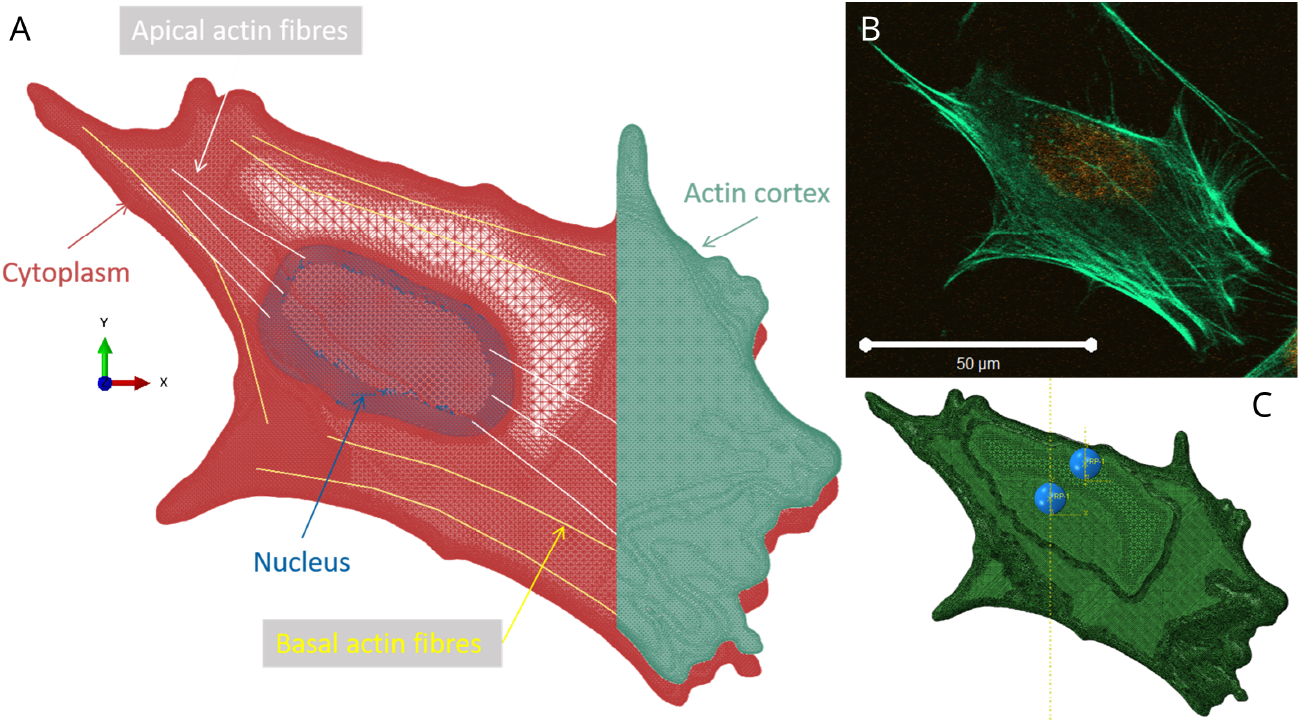
A: FE model for the single cell part, including nucleus (blue), cytoplasm (red), apical (white) and basal (yellow) actin bundles and actin cortex (green). The actin cortex covers the cell surface completely and it is in contact with it, the depicted representation was chosen for clarity only. B: The basal confocal section of MLO-A5 cells stained for actin (green) and nuclei (orange) is shown. Scale bar equal to 50 *μm*. Images are representative of about 5-6 cells imaged per slide; three slides were imaged. C: FE model comprising the cell and the indenting beads; the bead locations used for both indentation regions are shown for representation purposes only, as separate simulations were run for nuclear and peripheral indentations.

### Atomic force microscopy measurements

The acquisition protocol of the AFM data is described elsewhere [10]. Briefly, spherical tipped cantilevers were used to indent 180 spatially separated single cells over 3 separate experiments. Each cell was indented 15 times on the nucleus and the periphery using a 5-point 3 *μm*-spaced grid [16]. The obtained data files were analysed in MATLAB (Mathworks) to calculate the experimental effective modulus by Hertz model fitting [5,7]. Nanoindentation data were averaged to obtain a global effective stiffness for the nuclear and peripheral regions for each cell.

### Finite element modelling

The cell chosen for the model design represented the population morphology: its dimensions for the area, major and minor axes of the nucleus and the whole cell were in the ranges for the population average as measured in preliminary experiments (data not shown). The following steps were performed in order to create the single cell FE model:

1. Image segmentation: the confocal images were segmented in ImageJ [14] to obtain separate masks for the nucleus and cytoplasm.
2. Geometry reconstruction and meshing: segmented images were imported in Simpleware ScanIP (Synopsis, Inc.) to reconstruct three-dimensional volumes and to prepare the FE model by congruously meshing them with smoothed tetrahedral quadratic elements.
3. FE model refinement: the reconstructed geometry was imported as an input file in Abaqus (Simulia, Dassault Systèmes). Actin fibres and cortex were added, and boundary conditions and simulation parameters were set to simulate the indentation of a spherical bead at different locations over the cell.

#### Actin cytoskeleton modelling

Discrete truss elements were added representing the apical and basal actin bundles (Fig. 1A in white and yellow). Their organisation and dimensions were chosen from the confocal imaging information: the basal actin bundles roughly followed the morphology of the cell and were tied to the cytoplasm at nodes regarded as focal adhesions; the apical actin bundles were designed to join the nucleus and the cytoplasm in a parallel fashion [15, 18] and were therefore tied to the nucleus on one end. A cross-section of all truss elements was chosen equal to 0.2 *μm*^2^. The actin cortex (Fig. 1A in green) was modelled as a thin shell homogeneously covering the cell surface [1, 18]. A thickness of 0.2 *μm* was chosen [1, 8, 9]. The cell membrane was not included in the model as it was considered negligible in terms of mechanical deformation resistance, as it is much softer than the actin cortex [1, 18].

#### Material properties and simulations

Materials [18] and element types for each subcellular component are summarised in Table 1. Homogeneous, isotropic and elastic material properties were assumed. A hybrid formulation was attributed to all elements to account for the almost incompressible material behaviour. The spherical bead used as indenter was modelled as a rigid shell with radius equal to 3*μm* (Fig. 1C). The indentation was controlled by setting the bead displacement in the z-direction for 1 *μm*, with the ideal substrate on the xy-plane. The nodes at the cell base were constrained on the z-direction to model the cell adherent to the substrate. In the nodes denominated as focal adhesions, i.e. where the basal actin bundles were tied to the cytoplasm, an encastre condition was enforced. A contact interaction was defined between the bead and the top surface of the cell, with an isotropic friction coefficient of 0.001. A finite surface to surface sliding was allowed. Large deformation formulation was included [15]. Two separate versions of the model were run with the spherical bead indenting the cell nucleus and periphery, respectively, for comparison with the experimental data (Fig. 1C). To this aim, the total reaction forces in the z-direction were used against the bead displacement to plot force-displacement curves similar to the ones obtained with the AFM. An algorithm was written in MATLAB to fit these curves with the Hertz model and obtain the computational effective modulus. Simulations ran as batch jobs on high computing performance facilities (Iceberg, University of Sheffield) and took on average about one hour of system time when launched on a 2×8-core Intel E5-2670 node with 256 GB of RAM memory.

**Table 1.**
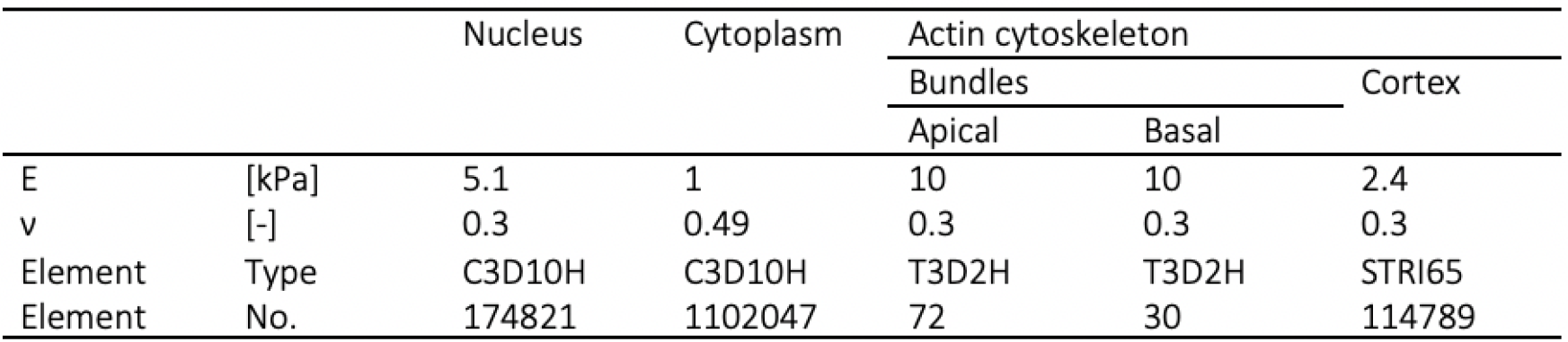
Material properties, Abaqus element type and number of elements of each modelled cellular component are summarised. An elastic material definition was used in terms of Young’s modulus (E) and Poisson’s ratio (*ν*).

## Results

A total of 2137 and 2556 indentations across 168 and 180 different MLO-A5 cells were analysed for the nucleus and periphery, respectively (Fig. 2). The median and interquartile range for the experimental effective modulus were calculated as 2.8 (1.5) *kPa* and 6.9 (4.7) *kPa*, for the nucleus and periphery. Two separate simulations were run to simulate AFM nanoindentation on the cell nucleus and periphery. When indenting the cell periphery, the simulation reached convergence and the total displacement of 1 *μm* was achieved; when indenting the nucleus, the simulation did not complete due to excessive element distortion (maximum indentation of 580 *nm*). This was expected due to the irregular geometry of the single cell model obtained from confocal images. The computational effective modulus was obtained by fitting the Hertz model on the displacement vs. reaction forces on the z-direction. The comparison between the FE and single cell AFM results for a sample of MLO-A5 tested over the nucleus and the periphery is shown in Fig. 3. The indentation depth vs. force curves were comparable for the nucleus, but lower forces in the FE model were exerted at similar indentation depths in the case of the periphery (Fig. 3A). In the case of indentation over the nucleus, the effective modulus tended to increase with the indentation depth, while a constant average stiffness was calculated for the experimental values (Fig. 3B). Conversely, the indentations towards the cell periphery converged to an effective modulus value of about 1.2 *kPa* constant for increasing indentation depths. This value was lower than the experimental stiffness values obtained when testing the cell periphery. All computational effective modulus values were in the ranges of dispersion for experimental ones. The values found in the nucleus and periphery simulated indentations were different, due to the contribution to the overall stiffness of the different materials and their local morphology. Conversely to what was observed in the experiments, the computational effective modulus was higher on the nucleus.

**Figure 2.**
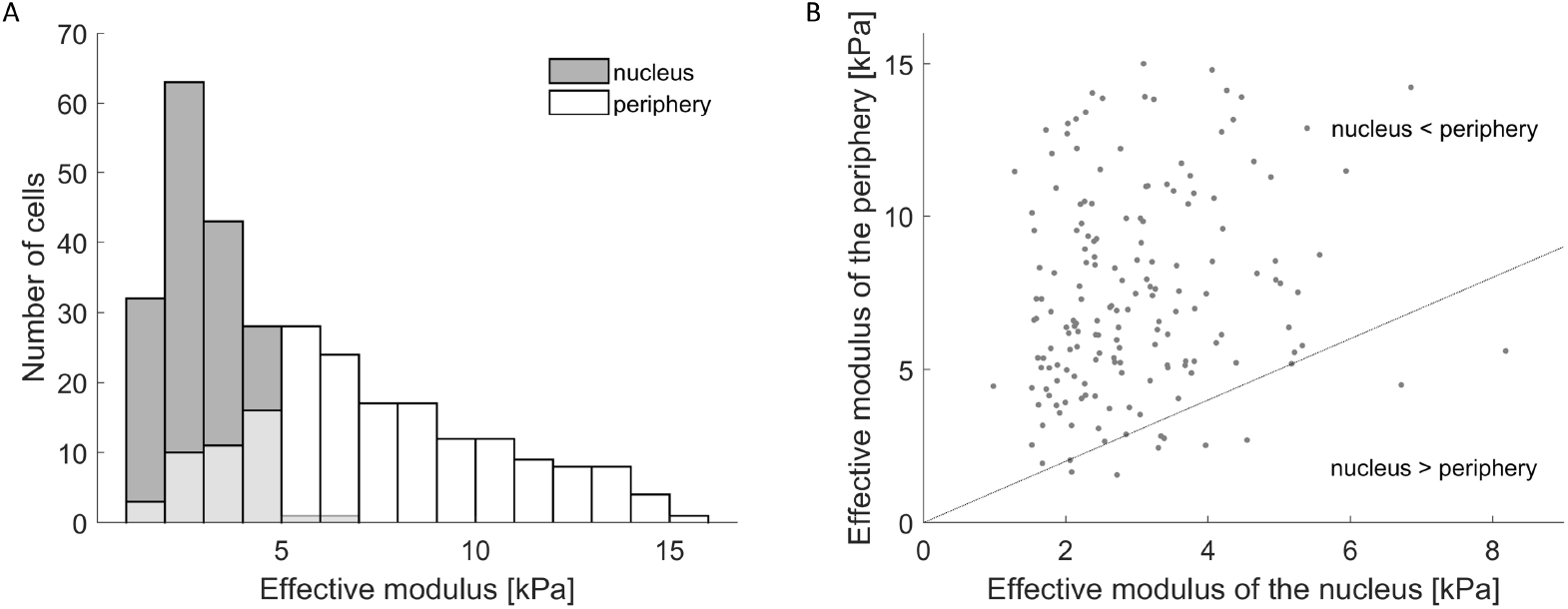
A: Histogram of single cell average effective moduli for MLO-A5 cells. Histograms for the nuclear area have shaded fill, histograms for the peripheral area have no fill. B: Comparison of the effective modulus of the nuclear and peripheral regions for each cell (dot marker).

**Figure 3.**
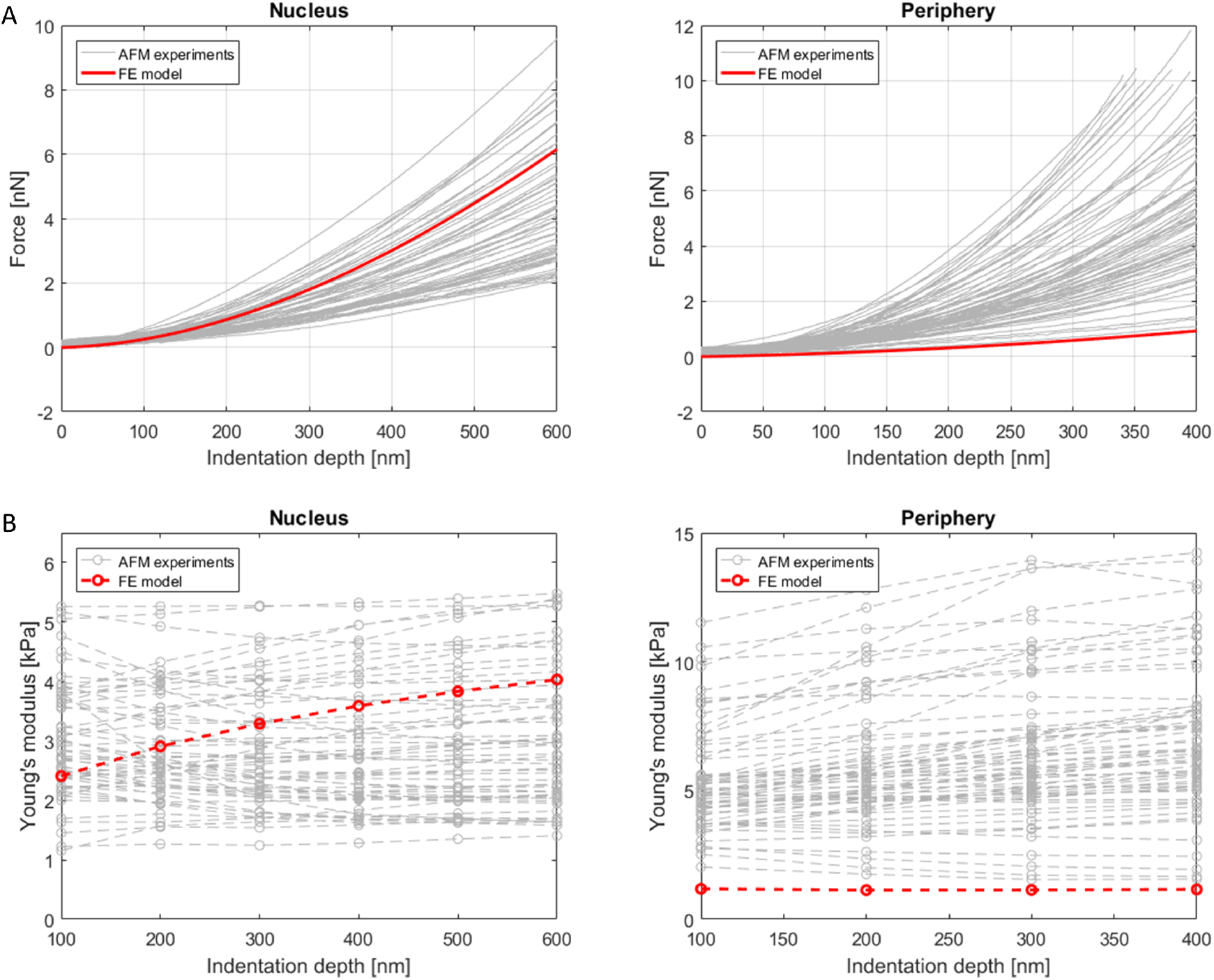
The FE results (red) were compared to the AFM experimental results for one sample of 80 MLO-A5 cells, for indentations over the nucleus (left) and the periphery (right). A: the indentation depth vs. force plots; B: the indentation depth vs. fitted effective modulus.

## Discussion

An FE model was created to simulate single cell indentation by AFM. The model included nucleus, cytoplasm, actin cortex and actin fibres, and was based on confocal images of a representative MLO-A5 cell. Simulations of indentation over the cell nucleus and periphery were performed, representing analogous AFM experiments. The computational effective modulus was different for the nucleus and periphery, highlighting the importance of the realistic model geometry and the inclusion of different cell components. The computational values were in the range of experimental values for MLO-A5 cells, despite showing higher stiffness for the nucleus than for the periphery, oppositely to the average experimental findings. However, it should be noted that some single cells also showed a similar behaviour experimentally (Fig. 2B). The cytoskeleton, and in particular actin, has been proposed as the major contributors to cell stiffness [9]. In the present model, the cytoskeleton was modelled only as the actin component, in the form of the actin cortex and bundles. The actin cortex played a role in withstanding the AFM bead compression and the bundles were involved in transmitting the load within the cell, as also suggested by experimental evidence [9]. However, the absence of other components could have impacted on the strain distribution, as it was suggested that myosin might reinforce the actin scaffold [4] and microtubules and intermediate filaments would be involved in force transmission [1, 2]. It has been previously shown that the spatial organisation of the actin fibres play a role in the overall cell stiffness, with the presence of aligned and/or peripheral actin fibres reinforcing the cytoskeleton [2,4]. The computational effective modulus obtained for the periphery was close to the lowest bound of dispersion of the experimental values, suggesting that the simplified modelling of the peripheral cytoskeleton caused the inability to capture this fibre reinforcement. It would be therefore of interest to model different single cells with known experimental effective moduli to evaluate the ability of the computational models to replicate experimental results. The FE material properties were obtained from different cell types [1, 18] and might not be accurate for MLO-A5 cells. The impact of changing the mechanical properties of single cell components on the overall simulated effective modulus has been previously investigated [2, 18]. The material properties could, therefore, be adjusted to obtain a better match to the average experimental values, however, this was not considered appropriate in this context not being able to back up any structural changes with experimental evidence. Another limitation could reside in the modelling of all materials as isotropic elastic, despite the fact that cells are heterogeneous in composition and present time-dependent mechanical properties [9]. However, similar hypotheses were considered when using the Hertz model and were shown accurate enough for the stated objectives [13]. Moreover, information on the time-dependent nature of single cell components material properties was not readily available in the literature, implying making modelling assumptions which would be hard to verify [2]. Finally, it should be mentioned that the computational model represented a static snapshot of the cell, as no dynamic polymerisation of the actin cytoskeleton was included and no molecular structures were taken into account [11]. The inclusion of different cell components and of a realistic cell morphology represented, however, an attempt towards more reliable single cell models and the importance of these features was highlighted.

## Acknowledgments

The work was supported by the EPSRC (“Multisim”, Grant No. EP/K03877X/1), the European Research Council (258321) and the University of Sheffield (“Mechanoreceptors in health and disease” Network Scholarship). The sponsors had no role in the study design, in the collection, analysis and interpretation of data; in the writing of the manuscript; and in the decision to submit the manuscript for publication. SM wishes to thank Dr N Gavara and Prof J Hobbs for useful and constructive discussion.

